# GABA–GABA_A_ Receptor Signaling Orchestrates Invasion and Metastasis in Triple Negative Breast Cancer

**DOI:** 10.1101/2025.10.03.674636

**Authors:** Esther Afolayan, Boxon Zhang, Mingde Han, Chloe A.K. White, Merlyn Emmanuel, Thomas J. Velenosi, Karla C. Williams

## Abstract

Cancer is a leading cause of death globally, with the majority of cancer-related deaths resulting from cancer metastasis -the process by which cancer cells disseminate to distant sites. To metastasize, cancer cells acquire traits in support of diverse cellular processes that enable dissemination, survival, and colonization. Tumor cell dissemination requires invasion at local and distant sites and this process can be influenced by intrinsic and extrinsic factors. Here, we investigate the role of the neurotransmitter gamma-aminobutyric acid (GABA) in triple-negative breast cancer (TNBC) invasion and metastasis. TNBC cells increased invasion in response to GABA and this was found to be mediated through the GABA_A_ receptor family. TNBC cell lines were found to be responsive to exogenous GABA and also produced endogenous GABA. Pharmacological inhibition of GABA_A_ receptors reduced TNBC invasion and cancer cell dissemination and resulted in inhibition of GSK3α activity. TNBC cell lines were found to express the GABRE subunit and loss of GABRE impaired GABA-mediated invasion and tumor cell dissemination. These findings support a role for GABA signaling through GABA_A_ receptors in mediating TNBC progression.

## INTRODUCTION

Breast cancer is the most commonly diagnosed malignancy among women worldwide and a leading cause of cancer-related mortality (1). Triple-negative breast cancer (TNBC), which lacks expression of estrogen receptor, progesterone receptor, and HER2, accounts for approximately 15–20% of breast cancer cases. TNBC is characterized as an aggressive cancer with high rates of metastasis, poor 5-year survival rates, and limited targeted treatment options, compared to other breast cancer molecular subtypes (2). Elucidating the molecular mechanisms driving invasion and metastatic dissemination in TNBC is critical both to deepening our understanding of metastasis and uncovering potential new therapeutic targets.

Emerging evidence has highlighted a role for neurotransmitters in cancer proliferation and progression (3,4). Gamma-aminobutyric acid (GABA) is a key inhibitory neurotransmitter in the central nervous system. GABA functions exclusively through GABA_A_ and GABA_B_ receptors (5,6). GABA_A_ receptors are ionotropic receptors that gate chloride channels, while GABA_B_ receptors are G-protein coupled receptors. These receptors can be differentiated on the basis of their antagonist selectivity. Selective and potent antagonists of GABA_A_ and GABA_B_ receptors (gabazine and bicuculline (7) and 2-Hydroxysaclofen,(8) respectively) support studies investigating the role of GABA and GABA receptor families in normal physiology and pathophysiology. Prior work from our group, and others, have identified a pro-invasive role for GABA in cancer (9-12). Breast cancer cell lines exposed to GABA increase the formation of invasive structures termed invadopodia and increase cancer cell extravasation,(9) and elevated levels of GABA have been reported in breast and colon tumors (13,14). In breast cancer clinical specimens, GABA_A_ receptor expression has been documented, with elevated expression in breast cancer brain metastasis relative to primary tumor tissue (15,16). These studies provide important insights into the role of GABA in breast cancer, but appreciably more work is needed in order to understand how GABA accumulates in tumors and the molecular mechanism underlying GABA-mediated breast cancer progression. Here, we investigate the role of GABA_A_ and GABA_B_ receptors in breast cancer and evaluate the cellular and molecular response of TNBC cell lines to GABA.

## RESULTS

### GABA functions through GABA_A_ receptors to regulate TNBC cell invasion

Previously, we demonstrated that invasive structures, termed invadopodia, promote breast cancer cell dissemination and respond to the addition of exogenous GABA which increased tumor cell extravasation. To further characterize the role of GABA in breast cancer, specifically TNBC, TNBC cell lines BT-549 and MDA-MB-231 were incubated with GABA (1mM) and rates of invadopodia formation and tumor cell invasion were assessed. Exogenous addition of GABA significantly increased the percentage of invadopodia-forming cells in both the BT-549 and MDA-MD-231 cell lines (Fig. 1A-D). Invadopodia can promote localized tumor cell invasion (17-19), through matrix degradation, and tumor cell dissemination, through transendothelial invasion across a vascular endothelium (18,20-22). To examine the role of GABA in stimulating invadopodia-mediated processes we evaluated tumor cell invasion. GABA-stimulated invasion through a matrix barrier was evaluated using a transwell invasion assay. The addition of GABA resulted in a significant increase in tumor cell invasion for both the BT-549 and MDA-MB-231 cell lines (Fig. 1E-F).

**Fig 1.**
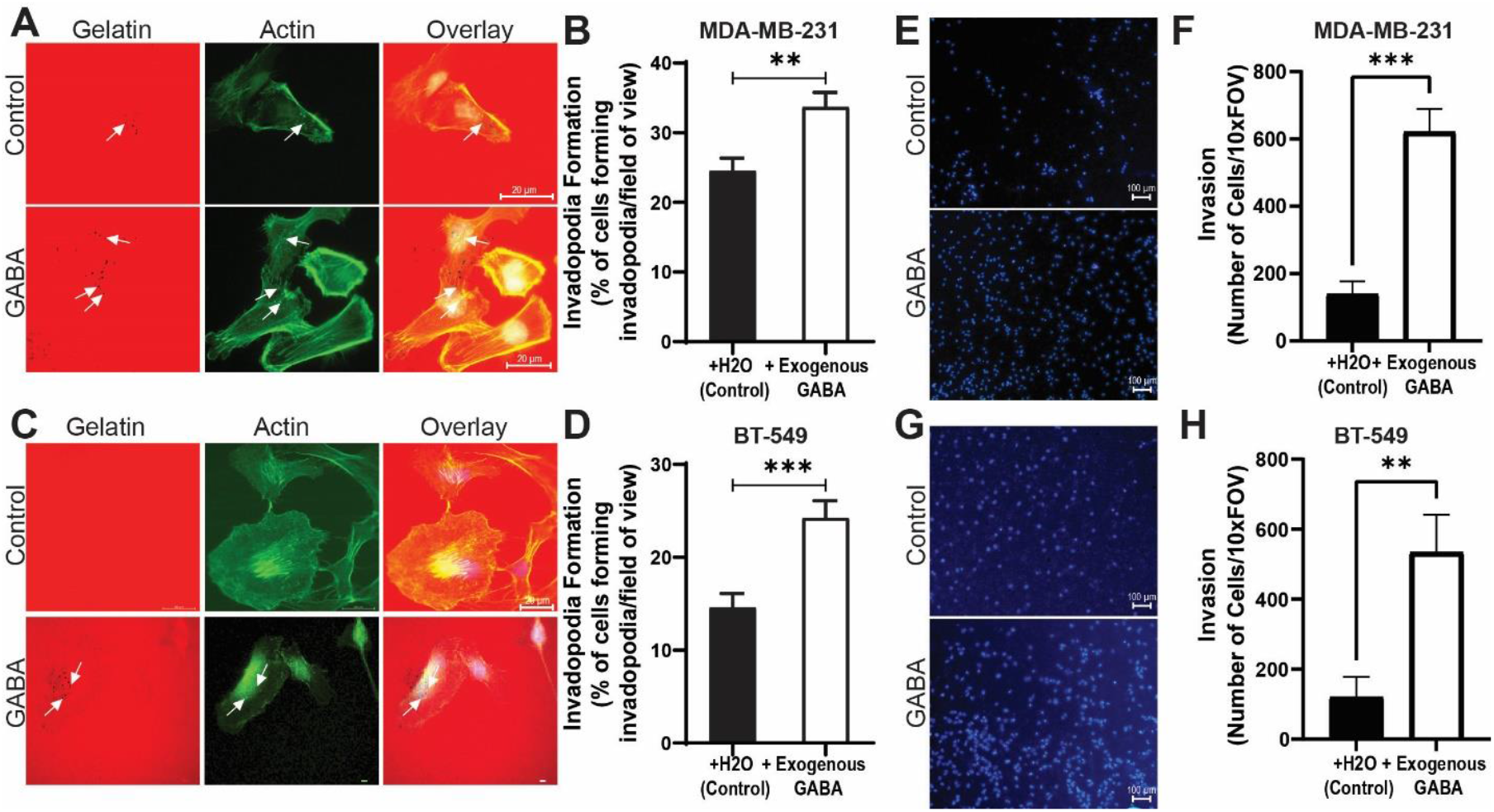
GABA promotes invadopodia formation and invasion in TNBC cell lines. (A, C) Representative images of invadopodia formation in MDA-MB-231 (A) and BT-549 (C) cell lines. Cells were plated on Alexa594-labeled gelatin-coated coverslips and incubated at 37 °C for 5h in the dark. Cells were fixed and stained with Alexa488-phalloidin to visualize F-actin. Images were acquired with a Zeiss Axio Observer microscope. White arrows indicate gelatin degradation mediated by invadopodia. (B, D) quantification of invadopodia formation rates in (B) MDA-MB-231 and (D) BT-549 cell lines. Quantification represents the percentage of invadopodia-positive cells from three independent experiments, with 20 randomly sampled fields per condition. Data show mean ± SEM. Student’s t-test. ^**^p < 0.01, ^***^p < 0.001. (E, G) Representative transwell invasion images and (F, H) quantification for MDA-MB-231 (E, F) and BT-549 (G, H) cells. Cells were serum-starved overnight, seeded (7×10^4^) into Matrigel-coated transwell inserts, and allowed to invade for 22 h toward serum-free medium ± GABA (1 mM). After fixation, the insert membrane was excised, and the lower surface was mounted with antifade mountant containing DAPI and imaged. Quantification reflects the mean number of invaded cells per field from three independent experiments (four randomly selected fields/insert). Graphs show mean ± SEM. Student’s t-test. ^**^p < 0.01, ^***^p < 0.001.

Having established a role for GABA in invadopodia formation and invasion, we sought to identify the GABA receptor family mediating GABA response in TNBC. To determine whether GABAergic regulation of invadopodia formation is mediated through GABA_A_ and/or GABA_B_ receptors, BT-549 and MDA-MB-231 cells were treated with the GABA_A_ receptor inhibitor, gabazine, or the GABA_B_ receptor inhibitor, 2-OH-saclofen. Both inhibitors were either administered alone or in combination with exogenous GABA. When co-administered with GABA, the percentage of invadopodia-forming cells was significantly lower in gabazine-treated BT-549 and MDA-MB-231 cells compared with 2-OH-saclofen–treated, relative to the control condition (Fig. 2A-B). This demonstrates that, in these cell lines, GABA acts through GABA_A_ receptor family members. Gabazine is a competitive inhibitor of GABA binding to the GABA_A_ receptor. Therefore, in the absence of exogenously added GABA, we hypothesized that gabazine treatment would not impact the rates of invadopodia formation. In contrast, we observed a significant reduction in invadopodia formation in gabazine-treated MDA-MB-231 cells relative to our controls. A trend in reduction was also found for the BT-549 cell line, but this did not meet significance. From this, we considered the possibility that TNBC cells produce GABA and that the observed reduction in invadopodia formation in the presence of gabazine likely stemmed from the inhibition of endogenous GABA binding to its cognate receptors. To test for endogenous GABA production, we analyzed GABA metabolite levels in BT-549 and MDA-MB-231 cell lines by LC-MS. GABA was found to be produced by both BT-549 and MDA-MB-231 cell lines (Fig. 2C).

**Fig 2.**
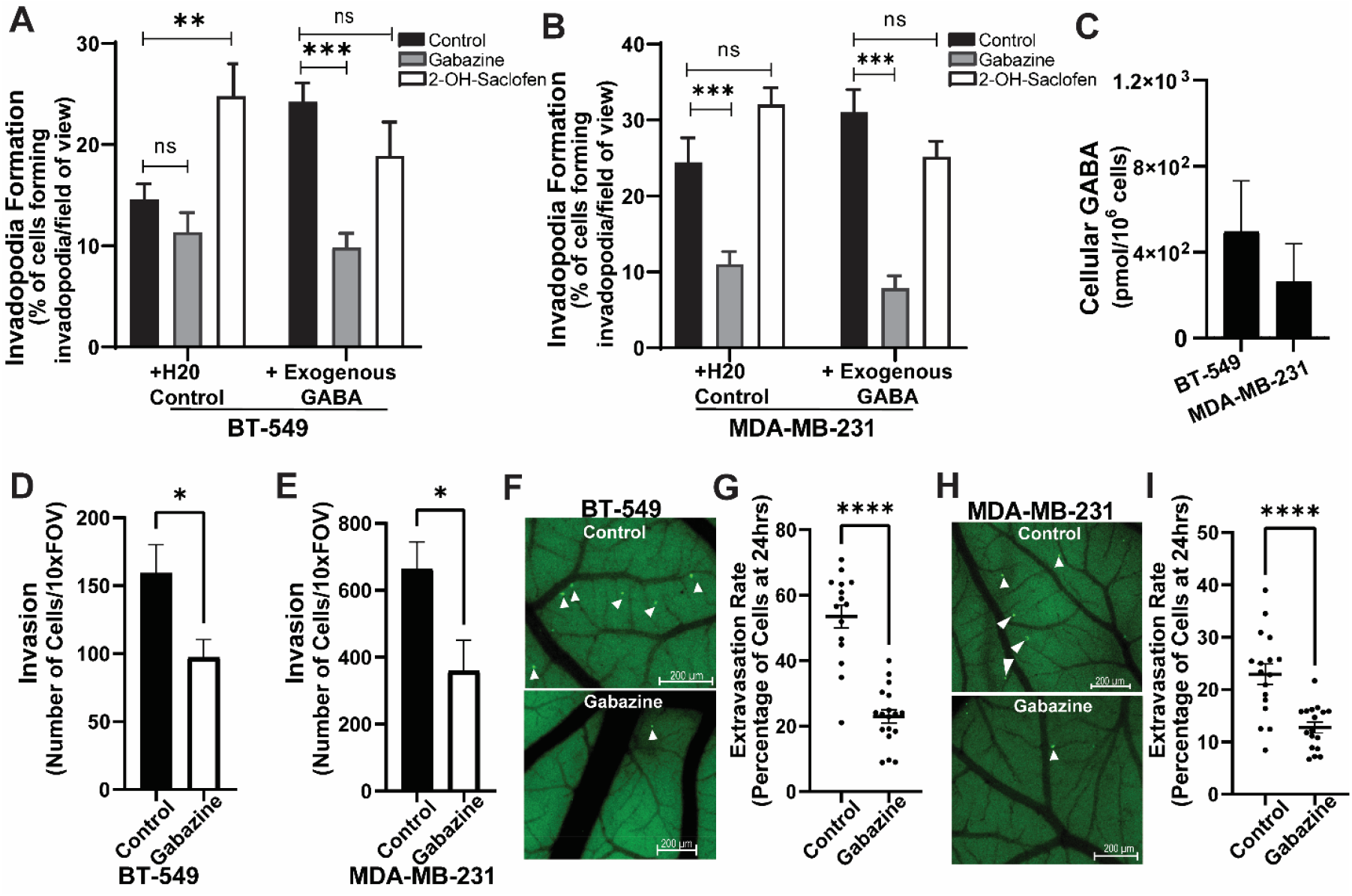
Pharmacological inhibition of GABA_A_R reduces invadopodia formation and reveals endogenous GABA production. (A, B) Quantification of invadopodia formation in BT-549 (A) and MDA-MB-231 (B) cells treated with GABA_A_R antagonists: gabazine (150 µM) or 2-OH-saclofen (300 µM), with or without GABA (1 mM). Data show mean ± SEM from three independent experiments, with 20 randomly sampled fields per condition. One-way ANOVA with Bonferroni post hoc test. ns: not significant, ^**^p < 0.01, ^***^p < 0.001. (C) LC–MS quantification of endogenous GABA in TNBC cell lysates. (D, E) Quantification of tranwell invasion of BT-594 (D) and MDA-MB-231 (E) cell lines. Cells were serum-starved overnight, seeded (7×10^4^) into Matrigel-coated transwell inserts along with either gabazine or vehicle control (water), then allowed to invade for 22 h toward serum-free medium with GABA (1 mM) ± gabazine (150 µM). After fixation, the insert membrane was excised, and the lower surface was mounted with antifade mountant containing DAPI and imaged. Quantification reflects the mean number of invaded cells per field from three independent experiments (four randomly selected fields/insert). (F-I) Cancer extravasation rates assessed using the chick embryo chorioallantoic membrane (CAM) model. BT-549 and MDA-MB-231 cell lines stably expressing ZsGreen were intravenously injected into the CAM. Cancer cells in the vasculature were quantified post-injection (T=0) and post-extravasation (24hrs post-injection). (F, H) Representative image of extravasated cancer cells in BT-549 (F) and MDA-MB-231 (I) cell lines (24hrs post-injection). (G, I) Quantification of extravasation rates in BT-549 (G) and MDA-MB-231 (I) cell lines. (D-I) Student’s t-test. Graphs show mean ± SEM. ^*^p < 0.05, Values are mean ± SEM, n = 3 biological replicates. ns: not significant, ^**^p < 0.01, ^***^p < 0.001.

2-OH-saclofen treatment showed no significant reduction in invadopodia formation. Consequently, to rule out the possibility that the observed lack of significance was due to low inhibitor concentration, both cell lines were treated with higher doses of 2-OH-saclofen (1 mM). No significant changes were observed in the percentage of invadopodia-forming cells (Supp. Fig. 1A). To further validate that GABA-stimulated invadopodia formation is mediated by GABA_A_ receptors, cells were treated with a second selective GABA_A_ receptor antagonist, bicuculline-methiodide. Bicuculline-methiodide produced similar results to those observed for gabazine treatment (Supp. Fig. 1B). Next, we evaluated whether GABA influences proliferation or migration using a wound healing assay. No significant differences were observed in either cell line in response to GABA treatment or inhibition of GABA_A_ receptor signaling (Supp. Fig. 2A-D), suggesting GABA-mediated effects are more specific to cancer cell invasion.

To further assess the impact of GABA_A_ receptor activity in mediating TNBC invasion, TNBC cell lines were treated with gabazine and cancer cell invasion was assessed. Gabazine treatment was found to significantly impair TNBC cell invasion (Fig. 2D and E). Next, to assess the role of GABA_A_ receptor activity in TNBC cell dissemination, an assessment of cancer cell movement out of blood vessels, extravasation, was performed. For this, we utilized the chick embryo chorioallantoic membrane (CAM) model as it is an established model for monitoring and quantifying cancer cell extravasation (9,21,23). Gabazine treated TNBC cell lines (ZsGreen-BT-549 and ZsGreen-MDA-MB-231) were intravenously injected into the CAM of the chick embryo and extravasation rates were quantified 24 h post-injection. Gabazine treatment significantly reduced TNBC cell extravasation compared to control treated cells (Fig. 2F-I). These results collectively support a model in which GABA promotes cancer cell invasion and dissemination in TNBC cells through GABA_A_ receptor activation.

### GABA_A_ receptor activity alters GSK3α activity

Our finding that GABA stimulates invadopodia formation through GABA_A_ receptors, prompted further mechanistic studies to identify downstream pathways activated by GABAergic signaling. GABA_A_ receptor signaling has been reported to stimulate ERK activity, specifically through the π subunit (24,25). GABA_B_ receptor signaling has been reported to regulate GSK3 activity, resulting in inhibitory phosphorylation of GSK3α/β (26). Given that PI3K/AKT signaling can regulate GSK3 activity and is frequently dysregulated in breast cancer, particularly in TNBC subtypes (27), we sought to investigate whether GABAergic-mediated regulation of invadopodia involves modulation of GSK3 activity. To test whether GABA_A_ receptors modulate AKT and GSK3 signaling, we assessed the phosphorylation state of AKT and GSK3α in BT-549 and MDA-MB-231 cells following GABA or gabazine treatment. AKT1 phosphorylation levels did not change with GABA or gabazine treatment (Supp. Fig. 3A,B). However, gabazine treatment significantly increased GSK3α (Ser-21) phosphorylation (Fig. 3A-D), suggesting that GSK3α is selectively responsive to GABAergic signaling and this is regulated through an alternative, non-AKT, pathway. For further evaluation of GSK3α activity in relation to invadopodia formation and mediating the response to GABA, we utilized the GSK3α selective inhibitor BRD0705 (28). Treatment with BRD0705 significantly reduced invadopodia formation in TNBC cell lines and impaired their response to exogenous GABA (Fig. 3E and F). This demonstrates that GSK3α activity is involved in invadopodia formation and suggests that GABA-mediated signaling through increased GSK3α activity regulates invadopodia formation.

**Fig 3.**
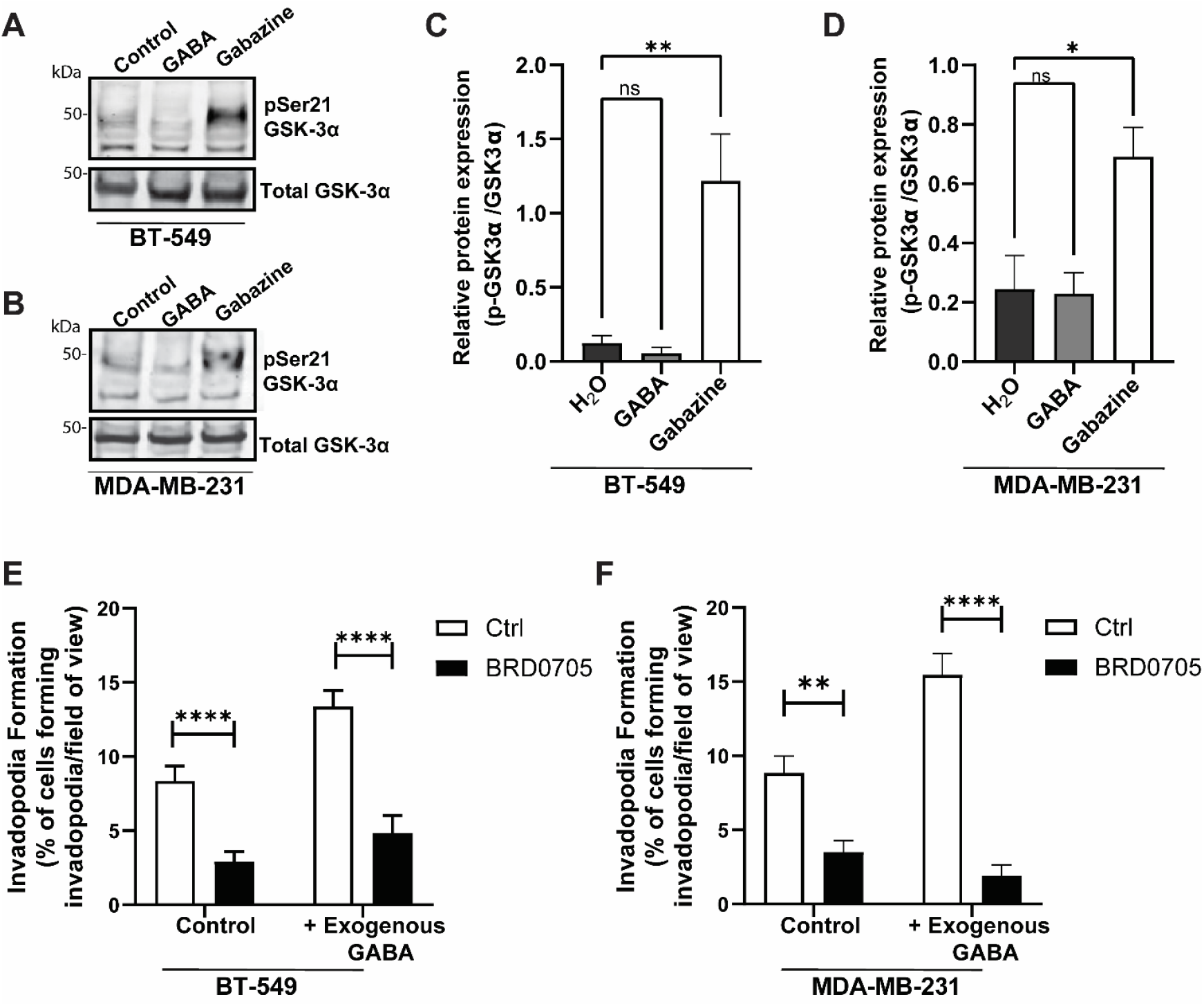
GABA_A_R signaling modulates GSK3α activity in TNBC cells. (A, B) Representative western blots of total GSK3α and phosphorylated GSK3α (Ser21) in BT-549 (A) and MDA-MB-231 (B) cell lines. Cell lines were treated with vehicle (H_2_O), GABA (1 mM), gabazine (150 µM, for 5 h. n = 3. (C, D) Quantification of GSK3α (Ser21) phosphorylation relative to total GSK3α in BT-549 (C) and MDA-MB-231 (D) cell lines. n = 3. Graphs show mean ± SEM. One-way ANOVA with Bonferroni post hoc test. (E, F) Quantification of invadopodia formation in BT-549 (E) and MDA-MB-231(F) cell lines treated with selective GSK3α inhibitor (BRD0705). Cell lines were plated on Alexa594-labeled gelatin-coated coverslips and incubated with GSK3α inhibitor, BRD0705 (10 µM) or vehicle (DMSO) ± GABA (1 mM). Quantification represents the percentage of invadopodia-positive cells from independent experiments, with 15 randomly sampled fields per condition. Data show mean ± SEM. One-way ANOVA with Bonferroni post hoc test. ^**^p < 0.01, ^***^p < 0.001, ^****^p < 0.0001.

### GABRE and GABRP expression are elevated in TNBC and expressed by TNBC cell lines

To identify the major GABA_A_ receptor subunit(s) expressed in TNBC, we analyzed GABA_A_ receptor subunit expression profiles in clinical breast cancer samples using the METABRIC database (29,30). Among the GABA_A_ receptor subunits, GABRD, GABRE and GABRP were among the most abundantly expressed subunits (Fig. 4A). GABRD, GABRE and GABRP were further evaluated and expression was grouped based on a classical three-gene identification system (Fig. 4B-D). GABRE and GABRP expression was found to be significantly higher in TNBC (ER-/HER2-) relative to all other breast cancer subtypes (Fig. 4C-D).

**Fig 4.**
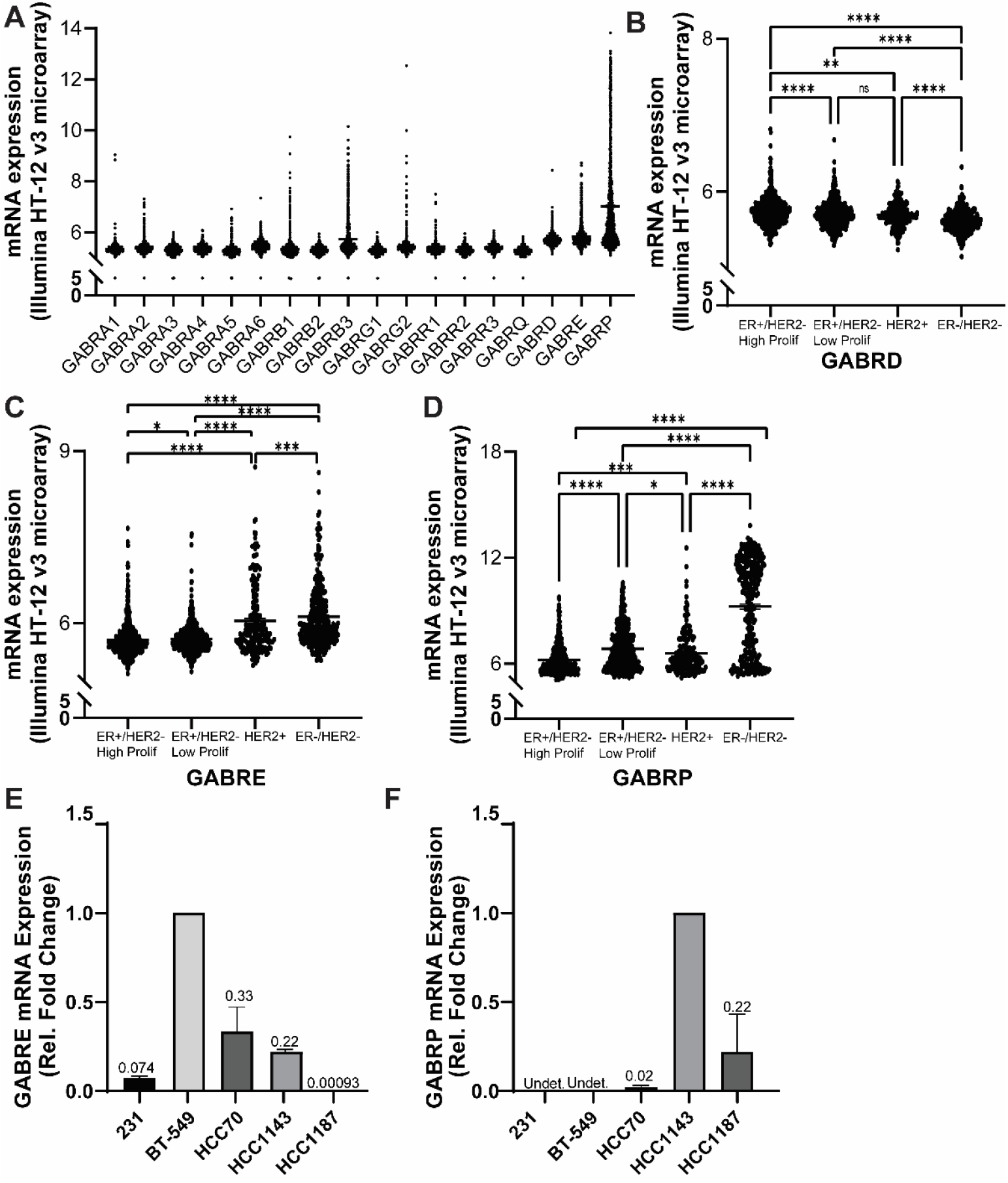
GABRE and GABRP are upregulated in TNBC, and GABRE is required for invadopodia formation. (A) METABRIC database analysis of GABA_A_R subunit expression in clinical breast cancer samples based on mRNA levels. (B–D) Expression of (B) GABRD, (C) GABRE, and (D) GABRP grouped by the classical 3-gene classifier. Data accessed via cBioPortal. Kruskal-Wallis test. ^*^p<0.05, ^**^p < 0.01, ^***^p < 0.001, ^****^p < 0.0001. (E, F) Quantitative RT–PCR validation of (E) GABRE and (F) GABRP expression in TNBC cell lines (n = 3).

Next, we assessed GABRE and GABRP mRNA expression in BT-594 and MDA-MB-231 cell lines. BT-549 and MDA-MB-231 cell lines expressed GABRE and lacked GABRP expression (Fig. 4E-F). As BT-549 and MDA-MB-231 cell lines have gene expression signatures similar to the claudin-low TNBC subtype (31), we also included additional TNBC cell lines classified as basal-like TNBC: HCC70, HCC1143, and HCC1187 (32). GABRP expression was found for the HCC1143, HCC1187, and HCC70 cell lines (Fig. 4F). HCC1143 and HCC70 were also found to express GABRE (Fig. 4E). Taken together, GABRE and GABRP are expressed in TNBC patient tumors and cell lines.

### GABRE promotes TNBC cell invasion and metastasis

Given our results demonstrating that the BT-549 and MDA-MB-231 cell lines expressed GABRE, but not GABRP, and that GABRE expression was the highest in BT-549, we selected the BT-549 cell line for further study. We hypothesized that GABA acts through GABRE to stimulate invadopodia formation, regulate GSK3α activity, and promote invasion. To assess this, BT-549 GABRE knockdown cell lines were generated (Fig. 5A). BT-549 control and GABRE knockdown cell lines were assessed for their ability to form invadopodia in the presence and absence of exogenously added GABA. Knockdown of GABRE significantly reduced invadopodia formation rates in the absence of exogenous GABA (Fig. 5B). Loss of GABRE also impaired the response of BT-549 cells to GABA demonstrating that GABA functions through GABRE to stimulate invadopodia formation (Fig. 5B). Next, we assessed the role of GABRE in cancer cell invasion and in the regulation of GSK3α activity. GABRE knockdown significantly reduced BT-549 invasion, as determined by transwell invasion assay (Fig. 5C and D). An assessment of GSK3α activity in BT-549 control and GABRE knockdown cell lines found a significant increase in the inhibitory phosphorylation of GSK3α (pSer21) in the GABRE knockdown cell line compared to the control (Fig. 5 E and F), demonstrating a reduction in GSK3α activity. Together, these findings indicate that GABA_A_ receptors mediate TNBC invasion and cell signaling, with the ε subunit playing a critical role in this process.

**Fig 5.**
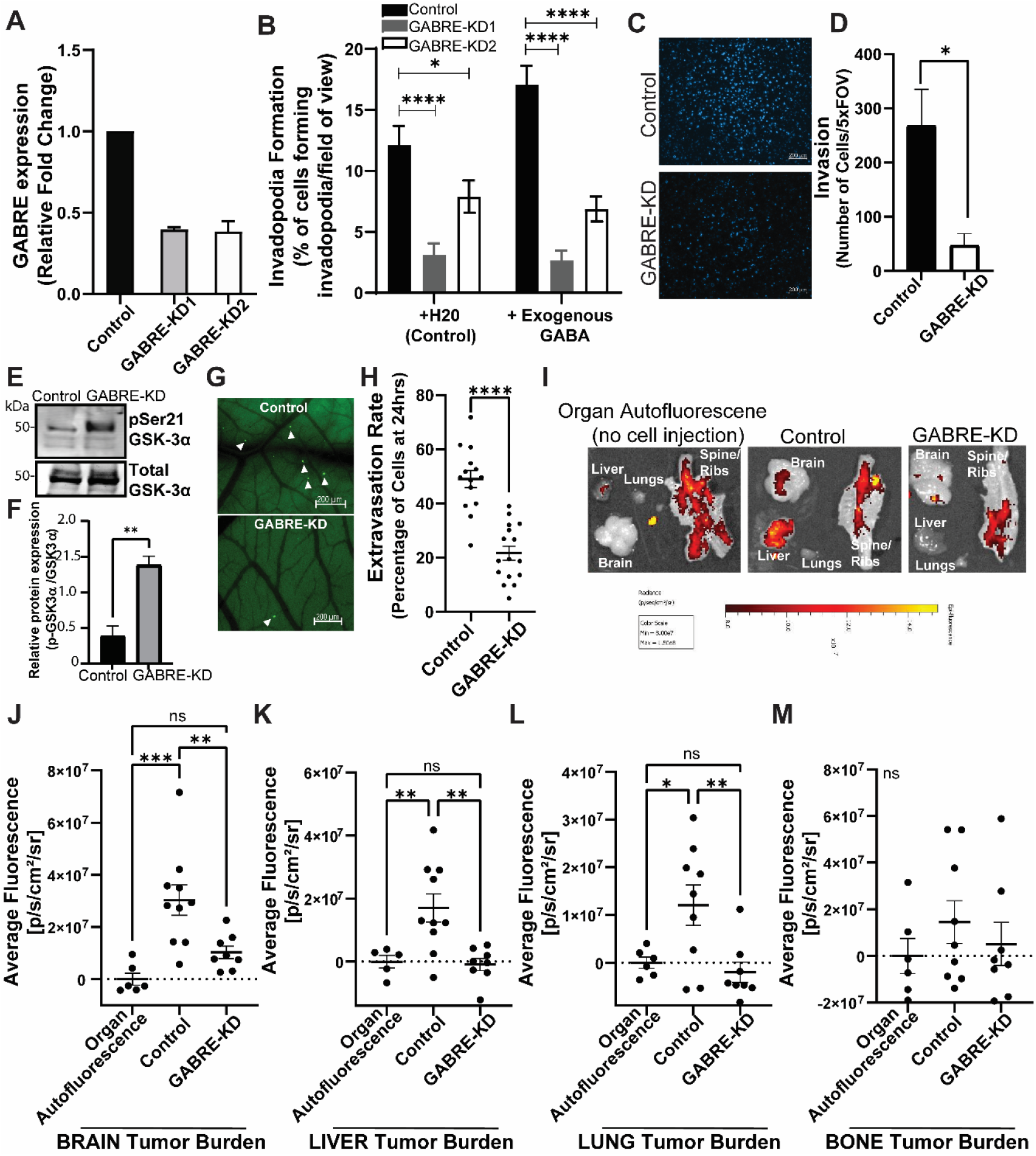
GABRE promotes TNBC cell line invasiveness both in vitro and in vivo. (A) Quantitative RT–PCR analysis of GABRE in BT-549 cells stably expressing shGABRE or scrambled (Scr) control. n = 3. (B) Quantification of invadopodia formation in BT-549 control and GABRE knockdown cell lines, with or without GABA (1 mM). Graphs show mean ± SEM. One-way ANOVA with Bonferroni post hoc test. n = 3. ^*^p < 0.05, ^****^p < 0.0001. (C) Representative transwell invasion images and (D) quantification for BT-549 control and GABRE knockdown cell lines. Cells were serum-starved overnight, seeded (7×10^4^) into Matrigel-coated transwell inserts, and allowed to invade for 22 h toward serum-free medium ± GABA (1 mM). Quantification reflects the mean number of invaded cells per field from three independent experiments. Graphs show mean ± SEM. Student’s t-test. ^*^p < 0.05. (E) A representative western blot of GSK3α (Ser21) phosphorylation and total GSK3α in BT-549 control and GABRE knockdown cell lines. (F) Quantification of GSK3α (Ser21) phosphorylation relative to total GSK3α in control and GABRE knockdown cell lines. (G,H) Cancer extravasation rates assessed using the chick embryo chorioallantoic membrane (CAM) model. BT-549 control and GABRE knockdown cell lines stably expressing ZsGreen were intravenously injected into the CAM. Cancer cells in the vasculature were quantified post-injection (T=0) and post-extravasation (24hrs post-injection). (G) Representative image of extravasated cancer cell lines (24hrs post-injection). (H) Quantification of extravasation rates. Student’s t-test. Graphs show mean ± SEM. ^***^p < 0.001. (I-M) Metastatic tumor burden assessed using the chick embryo model. BT-549 control and GABRE knockdown cell lines stably expressing ZsGreen were intravenously injected into the CAM. 7-days post-injection, organs were harvested and ex vivo imaged. (I) Representative images of tumor burden in organs from embryos injected with PBS alone, control cell line, or GABRE knockdown cell line. (J-M) Quantification of fluorescence signal in the brain (H), liver (I), lung (J) and bone (K). One-Way ANOVA with Fisher’s LSD test. Graphs show mean ± SEM. ns: not significant, ^*^p < 0.05, ^**^p < 0.01, ^***^p < 0.001.

To further ascertain the role of GABRE in TNBC cell invasion and cancer progression an evaluation of cancer cell extravasation and metastatic tumor burden was performed. Extravasation rates were evaluated for the ZsGreen BT-549 control and GABRE knockdown cell lines. Loss of GABRE was found to impair cancer cell extravasation and a significant reduction in extravasation rates was identified in the GABRE knockdown cell line (Fig. 5G and H). Using the chick embryo model, an evaluation of metastatic tumor burden was performed. The chick embryo model has been used to evaluate cancer cell dissemination and metastatic tumor burden across multiple cancer cell lines; it has also demonstrated utility in recapitulating metastatic organotropism in breast cancer cell lines (33,34). BT-549 control and GABRE knockdown cell lines were intravenously injected into the CAM. 7-days post-injection, organs were harvested and ex vivo imaging was performed. As a control, a subset of embryos was injected with PBS alone (i.e. no cancer cells) and organs were removed at end-point to establish the autofluorescence of each organ (Fig. 5I). Brain and liver metastases were consistently identified in embryos injected with the BT-549 ZsGreen control cell line (Fig. 5I-M). An evaluation of metastatic tumor burden in each organ identified a significant decrease in tumor burden in the brain, liver, and lung of embryos injected with the GABRE knockdown cell line (Fig. 5I-M). Collectively, these results point to a role for GABRE in promoting TNBC cell dissemination and metastasis.

## DISCUSSION

Here, we demonstrate that GABA signaling and GSK3α signaling converge in breast cancer, where GABA acts through GABA_A_Rs to promote invadopodia formation, tumor cell invasion, and metastasis, and selectively modulate GSK3α activity. GABA signaling has been implicated in diverse aspects of tumor biology, including cell proliferation, survival, and migration (9,12,14). In parallel, the serine/threonine protein kinase GSK3 is a key regulator of cytoskeletal dynamics, controlling actin and microtubule organization to support directional migration and invasion. Previous studies have demonstrated that GABA-mediated signaling regulates tumor progression in lung and colorectal cancers via GABA_B_ receptor regulation of GSK3β activity (11,26). Our results extend this paradigm by providing evidence of GABA-mediated signaling in the regulation or TNBC progression, via GABA_A_ receptors, with GSK3α serving as a key downstream effector in this pathway.

Although GABA primarily functions as the principal inhibitory neurotransmitter in the central nervous system, components of the GABAergic systems have been shown to be expressed in diverse cancer types, including breast cancer, and regulate tumor behavior (3,4). Our findings expand this literature by demonstrating that GABA functioning via GABA_A_ receptors, enhances TNBC invasion, and promotes metastatic dissemination. Notably, our findings reveal a selective expression of GABA_A_ receptor subunits across breast cancer subtypes. GABRE and GABRP were enriched in TNBC tumors. GABRE expression was found in claudin-low and basal-like TNBC cell lines, whereas GABRP expression was absent in claudin-low cell lines and mainly detected in basal-like cell lines. Our work focused on GABRE due to its expression in our cell lines of study (claudin-low: BT-549 and MDA-MB-231). Functionally, GABRE knockdown inhibited invadopodia formation and resulted in reduced cancer cell invasion in vitro and cancer progression in vivo, establishing a direct link between GABRE receptor subunit expression and TNBC metastasis.

The observed divergence between GABRE and GABRP expression patterns highlights the need to further consider the heterogeneity of GABA_A_ receptor composition in tumors and how it might govern distinct downstream signaling outputs. GABA_A_ receptors can functionally assemble in both homomeric and heteromeric forms (24,35), with receptor stoichiometry profoundly influencing ion channel kinetics, pharmacology, and downstream signaling specificity- factors which likely affect tumor progression (36,37). Future work should evaluate the role of GABRP in TNBC cell signaling and metastasis, and evaluate whether GABRE and/or GABRE function as a homomeric receptor or in heteromeric complexes.

Mechanistically, our results reveal that GABA_A_ receptor signaling converges on GSK3α, rather than AKT1 or GSK3β, to promote invadopodia formation. Although, GSK3 activity has been demonstrated to be modulated by GABAergic signaling in neurons via PI3K/Akt-dependent phosphorylation, our findings demonstrate a divergence from this canonical pathway (26). Our examination of the effects of pharmacological inhibition of GABA_A_ receptor activity, using gabazine, identified a significant increase in inhibitory GSK3α phosphorylation. The observed selectivity for GSK3α is particularly notable given the extensive focus on GSK3β in cancer biology. GSK3 is a highly conserved serine/threonine kinase that plays a central role in regulating several key cellular signaling pathways. GSK3α and GSK3β are highly homologous sequentially, with 97% sequence identity in their catalytic domain and overlapping substrates. Nonetheless, recent studies point to isoform-specific roles (38). Extensive studies have focused on GSK3β and its role in Wnt/β-catenin signaling (39). As GSK3β regulates β-catenin which acts as a tumor suppressor, GSK3β has been reported to play a tumor suppressive role in cancer. However, GSK3β can also regulate NF-κB signaling which has described oncogenic roles in cancer (40). The role of GSK3α in cancer is less well-described. Selective blocking of GSK3α activity using BRD0705 was shown to prolong survival in mouse models of acute myeloid leukemia; suggestive of a pro-oncogenic role for GSK3α (28). Our work extends this isoform-specific distinction to breast cancer, positioning GSK3α as a novel effector of GABA_A_ receptor signaling and TNBC invasion.

Together, our findings establish a mechanistic link between GABA_A_ receptor signaling and GSK3α activation in the regulation of TNBC cell invasion and metastasis. They also raise several important questions. In particular, the identity of the upstream kinase responsible for GSK3α phosphorylation remains unresolved. Additionally, it is unclear whether GABRE functions as part of a heteromeric receptor complex or signals independently as a homomeric receptor, an important distinction that could significantly influence downstream specificity. Furthermore, in vivo validation using mouse models would complement our ex ovo CAM assays and help clarify the relevance of this pathway to breast cancer progression. Finally, although our data highlights GABRE as a critical mediator in TNBC, GABRP may also play a distinct role in basal-like TNBC, representing a promising target for future investigation.

Finally, our results carry therapeutic implication. First, it raises the possibility that pharmacological targeting of select GABA_A_ receptors could disrupt TNBC invasion and perturb metastatic dissemination. Given that clinically approved GABA_A_ receptor modulators already exist, repurposing these drugs for oncology applications may be feasible. Second, our findings suggest that GABA_A_ receptor signaling regulates GSK3α specifically. Selective targeting of GSK3α, hence blocking the isoform-specific role of GSK3α, rather than broadly targeting both isoforms, may yield greater therapeutic precision and reduce unwanted effects.

## Acknowledgements

This work is supported by funding from the Canadian Institutes of Health Research (CIHR) Project Grant awarded to K.C.W. K.C.W. holds a Tier 2 Canada Research Chair.

## MATERIALS AND METHODS

### Cell Lines and Culture Conditions

MDA-MB-231 (ATCC; Cat. no.: HTB-26) cells were obtained from American Type Cell Collection (Gaithersburg, ML) and grown in Dulbecco’s Modified Eagle Medium (DMEM; Cat. No.: 319-005-CL, Wisent Bioproducts) supplemented with 10% fetal bovine serum (FBS; Cat. No.: 12483-020, Thermo Fisher Scientific). HCC-70 (ATCC; Cat. no.: CRL-2315), HCC-1143 cells (ATCC; Cat. no.: CRL-2321) and HCC-1187 cells (ATCC; Cat. no.: CRL-2322) were grown in Roswell Park Institute Medium (RPMI; Cat. No.: 350-007-CL, Wisent Bioproducts) supplemented with 10% FBS. The BT-549 cell line (ATCC; Cat. no.: HTB-122) was maintained in RPMI supplemented with 10% FBS and insulin (0.023 U/ml; Cat. No.: 12585-014, Gibco Life Technologies).

### Invadopodia Formation and Degradation Analysis

#### Preparation of Gelatin-Coated Sterile Coverslips

Square 22×22 mm glass coverslips were treated with 0.1% NaOH for 5 mins then washed twice with Phosphate-Buffered saline 1X (PBS; Cat. No.: 3112-425-CL, Wisent) before sterilizing in 70% ethanol for 5 mins. Next, coverslips were coated with 50 µg/ml poly-L-lysine for 20min (Cat. No.: 25988-63-0, Sigma), followed by 0.5% glutaraldehyde for 15min (Cat. No.: 111-30-8, Sigma) and then inverted on a 100 µl drop of Alexa594-conjugated gelatin for 10min. Labeled coverslips were then gently washed 3-times with PBS, incubated with non-essential amino acids (Cat. No.: 321-011-EL, Wisent) and washed extensively with PBS to remove excess reagents. The prepared coverslips were used immediately or stored in PBS at 4 °C till needed.

#### Invadopodia Degradation Assay

Cells were plated (1.2 - 1.5 × 105 cells/well) onto the prepared gelatin-coated coverslips with 1% penicillin/streptomycin (Cat. No.: P0781, Sigma) in 6-well plates. GABA (1 mM; Cat. No.: 56-12-2, Abcam), 2-OH-saclofen (300 µM - 1 mM; Cat. No.: 117354-64-0, Sigma), Gabazine (150 µM; Cat. No.: 104104-50-9, Abcam) or (-)-Bicuculline-methiodide (300 µM; Cat. No.: 40709-69-1, Abcam) were added, as indicated, and cells were incubated on coverslips +/-treatment for 5-6 hrs. At endpoint, cells were fixed with 4 % paraformaldehyde (PFA) for 8-10 minutes, gently washed twice with PBS, subsequently followed by immunofluorescence staining or stored overnight at 4°C in 0.1 M glycine (Cat. No.: 56-40-6, Thermo Fisher Scientific).

#### Immunofluorescence Staining

Fixed cells on coverglass were incubated in 0.1M glycine for 10-15min. Glycine was discarded from each well and cells were permeabilized with 0.1 % Triton X-100 for 2 min, then washed twice with PBS to remove excess reagents. Next, cells were blocked at room temperature (RT) in 5 % skim milk diluted in PBS for 1h 30 min, then washed twice with PBS before staining for F-Actin to detect invadopodia formation. Cells were blocked in 5 % skim milk diluted in PBS containing Phalloidin 488 (1:3000, Cat No.: A12379, Thermo Fisher Scientific) for 1h at RT were used to visualize F-Actin.

#### Quantifying Percentage of Invadopodia Formation

To quantify percentage of invadopodia formation, coverslips were imaged using a 63 ×/1.4 NA objective lens on a Zeiss Axio Observer fluorescence microscope (Carl Zeiss Inc., Heidelberg, Germany). Cells were classified as positive for invadopodia formation if gelatin spots of degradation (black holes) could be visualized under the cells. Cells were quantified as either invadopodia forming or non-forming in 20 fields of view randomly dispersed across the full surface area of each coverslip.

### Transwell migration and invasion assays

Transwell invasion assays were conducted using 24-well inserts with polyethylene terephthalate membranes (8.0 μm, Corning, NY, USA) pre-coated with matrigel (Corning). Cells were serum starved overnight prior to seeding. 100 μL cell suspensions containing 7 × 10^4^ cells were added to the upper chamber of the insert. Next, 600 μL DMEM medium containing 0.1% BSA and 1X PenStrep was placed into the bottom chamber, and either water or 1mM GABA added as a chemoattractant. 22 h later, non-invading cells on the upper surface of the membrane were removed. The inserts were gently washed twice with PBS, and the invaded cells on the lower surface were fixed with 4% PFA. Membranes were then excised, mounted onto glass slides using ProLong™ Gold Antifade Mountant with DAPI (Cat No.: P36935, Thermo Fisher Scientific), and imaged under a Zeiss Axio Observer fluorescence microscope. Four random fields per insert were selected for quantification. All experiments were performed in triplicate, and data are presented as mean ± SD.

### Live-Cell Proliferation Assay

Cell proliferation was monitored using the Incucyte Live-Cell Analysis System (Sartorius). BT-549 and MDA-MB-231 cells were seeded into 96-well flat-bottom tissue culture plates at a density of 2,500 or 10,000 cells per well, respectively, in 100 μL of complete growth medium supplemented with water, GABA and/or gabazine or 2-OH-Saclofen. Cells were allowed to adhere for 30 minutes to ensure uniform cell distribution before transferring into the Incucyte for imaging. Time-lapse imaging was initiated immediately, with scans acquired every 3 hours using phase-contrast optics. Cell proliferation was quantified by measuring percent confluence over time using Incucyte analysis software. Data were collected from triplicate wells and are presented as mean ± SD

### RNA Extraction

Cell culture samples were trypsinized then collected by centrifugation at 300 x g for 5 minutes at RT. Collected pellets were washed twice with PBS and centrifuged between washes then PBS was completely removed from the pellet. Subsequent steps were performed in a designated RNase-free environment; work surfaces and pipettes were cleaned with RNaseZap (Cat No.: R2020, Sigma) before commencing RNA extraction. GENEzol™ TriRNA Pure Kit (Cat. No.: GZXD100, Geneaid) was used to extract the total RNA from cell pellets according to the manufacturer’s instructions.

Cell pellets were lysed in 700 µL of GENEzol™ reagent by repeatedly pipetting solution till homogenized then left to sit a RT for 5 mins. Then 700 µL of anhydrous ethanol was added to each sample before vortexing and transferring solution to the supplied RNA-binding spin columns. Columns were centrifuged at 16,000 x g for 30 seconds in a benchtop centrifuge, treated with DNase I to remove any genomic DNA then washed and dried according to the manufacturer’s instructions. Finally, the purified RNA was eluted into 25-50 µL of RNase-free water (provided with the kit) and quantified at 260 nm on a spectrophotometer (Nanodrop, ThermoFisher Scientific). The purified RNA was stored at -80 °C till needed.

### Quantitative real-time PCR

Pre-designed, gene-specific TaqMan® probe and primer sets for GABRE (Cat. no.: 4331182, TaqMan® Gene Expression Assays, ThermoFisher Scientific) were used for gene expression analysis. The relative expression level of each gene was normalized to an endogenous control, GAPDH gene (Cat. no.: 4331182, ThermoFisher Scientific). For GABRP, predesigned GABRP primers and reference GAPDH primers were purchased from IDT (Cat. no.: NM_014211 and NM_002046, PrimeTime® Predesigned qPCR Assays, Integrated DNA Technologies) for use with the TaqMan probe. The relative expression of each gene was calculated using the 2-ΔΔCt method.

### Generation of control and GABRE knockdown cell lines

HEK 293T cells were seeded and grown overnight to approximately 70% confluence at the time of transfection. Lentiviral shRNA plasmids (scrambled control or GABRE-targeting; ABM, BC, Canada) together with packaging vectors PAX2 and MD2.G were transfected into 293T cells using Lipofectamine 3000 following standard procedures. Viral particles were collected 48 hours post-transfection before storing at −80℃. For infection, lentiviral supernatant was added to cultured cells with 8 μg/mL polybrene (Sigma-Aldrich). After incubation for 24 hours, infected cells were selected for 3 to 5 days with puromycin (Wisent).

### LC–MS quantification of endogenous GABA

Endogenous GABA levels were quantified using LC–MS following a previously described protocol (41). Briefly, cells were washed twice with PBS and harvested with TrypLE (Fisher Scientific, cat no. 13604013). After harvesting cells were counted, centrifuged at 500xg at 4℃ for 5 minutes, washed twice more with PBS, and flash frozen in liquid nitrogen. For sample preparation, cells were diluted in 50μL of 6% TCA containing 1 μM aminopimelic acid as the internal standard. Cells were sonicated for 10 minutes in a bath sonicator. All samples were centrifuged 15000 x g for 10 minutes and 10μL of the supernatant was transferred to a new tube containing 70μL of 100mM Borax (pH=9) with 1mM ascorbic acid, 1mM TCEP, and 100μM NaOH. Samples were derivatized with 20μL of 6-aminoquinolyl-N-hydroxysuccinimidyl carbamate (AQC, Cayman Chemicals, Item No. 30877) followed by a 10-minute incubation at room temperature and a 10-minute incubation at 55℃ to quench the reaction. Derivatized samples were diluted 10-fold in ammonium formate then injected onto a Waters Acquity UPLC BEH Phenyl Column (1.7μm, 2.1 x 50mm) at 40℃ with a flow rate of 0.6 mL/min in an ExionLC Series UPLC. The two mobile phases consist of 0.1% formic acid in water (A), and 0.1% formic acid in acetonitrile (B). Eluting metabolites were measured using a Sciex 7500 Triple Quad MS using the following transitions: GABA 274.1 > 171 m/z and aminopimelic acid 346.1 > 171 m/z.

### Immunoblotting

Cells were lysed in cold RIPA lysis buffer by scraping. Cell lysates (5-20 μg) were resolved on 4-12% bis-tris gradient gel by SDS-polyacrylamide gel electrophoresis and transferred onto polyvinylidene difluoride (Cat. no.: IPVH00010, Millipore) or nitrocellulose membranes (Cat. no.: A29911551, BioRad). Membranes were blocked in 5% skim milk OR 5% BSA for 1 hour and incubated with the following antibodies overnight at 4°C, phospho-GSK3α (1:10,000; Cat. no.: PA5-121291, Invitrogen), GSK3α (1:1,000; Cat. no.: MA5-15301, Invitrogen), and Vinculin (1:2,000; Cat. No.: V9131, Sigma), phospho-AKT1 (1:20,000; Cat. no.: 44-621G, Invitrogen), total AKT1 (1:1,000; Invitrogen). The appropriate HRP-conjugated secondary antibodies (1:3,000; Cat. no.: NA931V and NA934V, Amersham) were added to the membranes for 1 h.

Finally, the membranes were visualized using Immobilon Forte Western HRP substrate (Cat. no.: WBLUF0100, Millipore).

### Ex ovo chick embryo model

Tumor cell extravasation assay was performed using the chick chorioallantoic membrane (CAM) model was performed as previously described(23). Briefly, cell lines stably expressing ZsGreen were dissociated with trypsin, counted, washed with PBS and resuspended to 0.5 × 10^6^/ml, and 200 μl of the cell suspension was injected intravenously into each embryo. Post-injection, a 1/2inch grid was placed on the surface of the CAM cells and cells in the vasculature were counted. Twenty-four hours post-injection, extravasated cells in the marked region were counted and extravasation rates were quantified.

The tumor metastasis assay using the chick CAM model was performed as previously describe(33). Briefly, BT-594 cell lines stably expressing ZsGreen were dissociated with trypsin and counted. After washing three times with PBS, cells were resuspended to 0.5 × 10^6^/ml, and 200 μl of the cell suspension, was injected intravenously into each embryo. Seven days post-injection, embryos were sacrificed and organs were removed. A subset of embryos were injected with PBS alone as a control. Organs were imaged with IVIS Lumina (Caliper Life Sciences, Waltham, MA) using the fluorescence optical imaging setting for 30 sec. A region of interest (ROI) was drawn around each organ, and average fluorescence was measured for each ROI.

### Statistical Analysis

All data are expressed as mean ± SEM. All experiments were performed as, at minimum three, independent experiments. Data was handled in Microsoft Excel 16.90.2 and analyzed using GraphPad Prism 10.6.0. The normal distribution of the data was analyzed by Kolmogorov– Smirnov and Shapiro Wilk test. Normally distributed data were analyzed using an unpaired t test or Welch’s test. For comparison of two groups or more a one-way ANOVA followed by Bonferroni post hoc test was performed. For the chick embryo organ metastasis, the average signal obtain from all no-cell-injection control organ was subtracted from each corresponding organ of the control, GABRE-KD, and no-cell injection groups. from all the he P-value was obtained using GraphPad Prism. ^*^p < 0.05. ^**^p < 0.01. ^***^p < 0.001. ^****^p < 0.0001.

