## Supplemental Figure for "GABA–GABA_A_ Receptor Signaling Orchestrates Invasion and Metastasis in Triple Negative Breast Cancer"

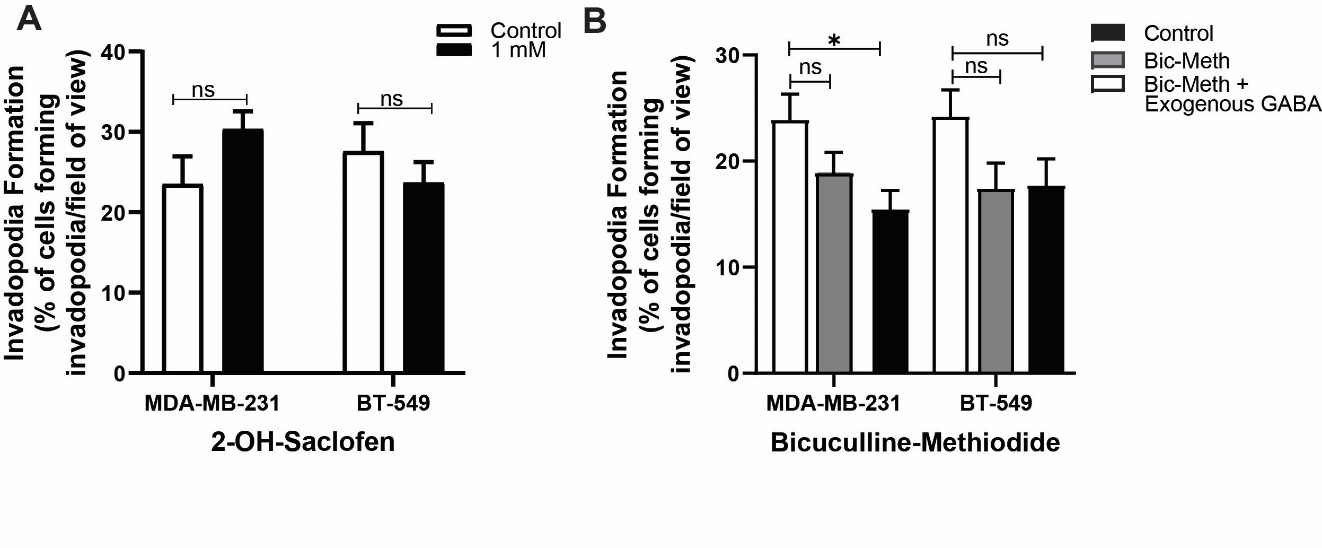

**Fig S1. GABA_A_R inhibitors suppress invadopodia formation.** (A) Quantification of invadopodia formation in MDA-MB-231 and BT-549 cells treated with 2-OH-saclofen (1 mM). (B) Quantification of invadopodia formation in cells treated with bicuculline-methiodide (300 µM). Graphs show mean ± SEM, n = 3. One-way ANOVA with Bonferroni post hoc test. ns: not significant, *p < 0.05.

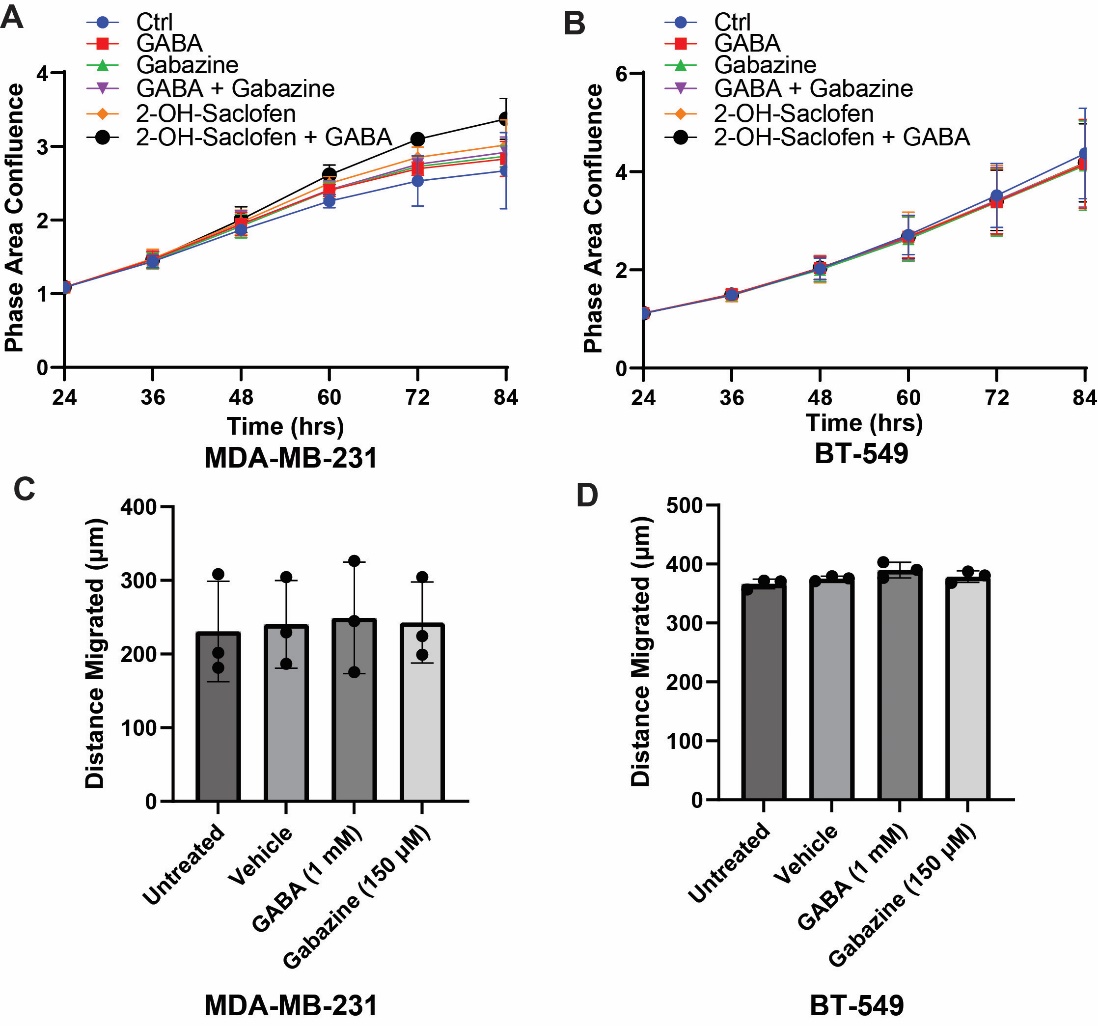

**Fig S2. Evaluating the Impact of GABA Signaling on TNBC Cell Proliferation and Migration.**
 (A, B) Proliferation rates of (A) MDA-MB-231 and (B) BT-549 cells treated with gabazine (150 µM) or 2-OH-saclofen (300 µM), with or without GABA (1 mM), assessed using the Incucyte live-cell imaging system. Data are mean ± SEM, n = 3. (C, D) Migration assays in (C) MDA-MB-231 and (D) BT-549 cells treated with GABA or gabazine. Migration rates were assessed using a wound healing assay, where cells were plated in a 96-well format and monitored via live-cell imaging on the Incucyte. Data are mean ± SEM, n = 3.

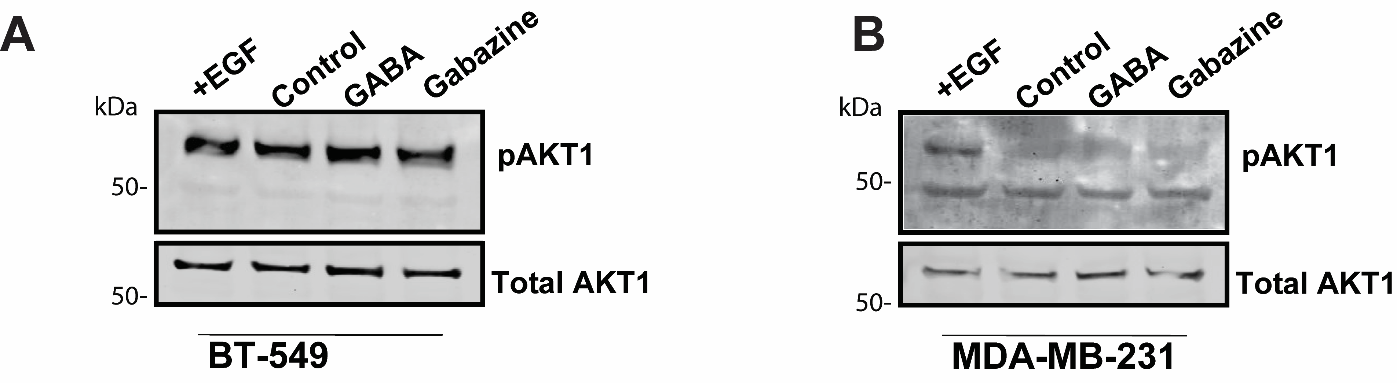

**Supp. Figure 3: AKT phosphorylation in TNBC cell lines is not regulated by GABA.** **(A, B)** Representative western blots of total AKT1 and phosphorylated AKT1 in BT-549 **(A)** and MDA-MB-231 cell lines **(B)**. Cell lines were treated with EGF (15min), or vehicle (H₂O), GABA (1 mM), gabazine (150 µM, for 5 h).
